# Genomic approach for conservation and the sustainable management of endangered species of the Amazon

**DOI:** 10.1101/2020.09.18.303115

**Authors:** Paola Fazzi-Gomes, Jonas Aguiar, Gleyce Fonseca Cabral, Diego Marques, Helber Palheta, Fabiano Moreira, Marilia Rodrigues, Renata Cavalcante, Jorge Souza, Caio Silva, Igor Hamoy, Sidney Santos

## Abstract

A broad panel of potentially amplifiable microsatellite loci and a multiplex system were developed for the Amazonian symbol fish species Arapaima gigas, which is currently in high danger of extinction due to the disorderly fishing exploitation. Several factors have contributed to the increase of this threat, among which we highlight the lack of genetic information about the structure and taxonomic status of the species, as well as the lack of accurate tools for evaluation of the effectivity of current management programs. Based on Arapaima gigas’ whole genome, available at the NCBI database (ID: 12404), a total of 95,098 unique perfect microsatellites were identified, including their proposed primers. From this panel, a multiplex system containing 12 tetranucleotide microsatellite markers was validated. These tools are valuable for research in as many areas as bioinformatics, ecology, genetics, evolution and comparative studies, since they are able to provide more accurate information for fishing management, conservation of wild populations and genetic management of aquaculture.

## 1. Introduction

The species *Arapaima gigas* (Schinz, 1822) belongs to the Arapaimidae family – order of the Osteoglossiformes [1], which composes one of the oldest groups of teleost fishes. It is the world’s largest scale fish, and specimens may reach up to 200 kg of body mass and 3 m of length [2]. It is an endemic species of the Amazon basin, being found in South American countries, such as Brazil, Peru, Colombia, Ecuador, Bolivia and Guyana [3,4]. In recent years, *A. gigas* has also received prominence in aquaculture due to its fast growth, high fillet yield, mild-flavored white meat, and great market acceptance, both domestically and abroad [5].

*A. gigas* is listed in Annex II of the International Convention on the Trade in Endangered Species (CITES) and in the International Union for Conservation of Nature’s (IUCN) Red List of Endangered Species, within the category “data deficient”, meaning that there is no proper information to make a direct or indirect assessment of its extinction risk based on its distribution and/or population status. Furthermore, in Brazil, arapaima were not included in the national list of endangered species due to lack of data [6].

Many reasons have contributed to the aggravation of the threat to *A. gigas*, including (i) the exploitation background of wild individuals, coupled with poor management strategies to comply with current fishing legislation; (ii) the current poorly refined genetic data on taxonomic structure and status; and (iii) the lack of management in certain areas, as well as the lack of accurate tools for evaluating the effectiveness on current management programs [7,8,9,10,11].

In Brazil, the predatory fishing of *A. gigas* – a practice which was intensified in the 1970s, impacted its population distribution in most parts of the Amazon basin. Consequently, strategies for management of the species were adopted by government agencies, such as the establishment of a minimum size for capture, the determination of reproductive season, and the annual fishing prohibition in the states of Amazonas, Pará, Amapá, Rondônia, Roraima, Tocantins and Acre [12,13,14]. However, the supervision of these strategies is deficient due to the lack of financial and human resources, as well as the lack of well-defined methods to identify the origins of the *A gigas*’s derived products, hindering the actions of the management system. For instance, recent data shows that 77% of *A. gigas* meat marketed in Santarém-PA comes from illegal fishing [15]. Furthermore, a substantial amount of *A. gigas* fresh or salty-conserved meat is exported illegally to Brazilian markets by middlemen, through the Guayaramerín border, in Bolivia [16].

The population structure of *A. gigas* is still undefined, and there are evidences showing the existence of more than one species of the genus Arapaima. On the other hand, the translocation of individuals is also an imminent threat [17], which can lead to loss of genetic diversity, decreased fitness and increased risk of extinction [18], since young individuals have often been translocated among different parts of the Amazon basin – and among this basin and others, mainly to attend aquaculture demand [19]. Therefore, the collection of molecular data is essential for monitoring and defining adequate strategies for the management actions.

It is a consensus that *A. gigas* populations are following a declining trend, where the main threat is overfishing [20,15], and the majority of communities do not practice sustainable management – with few exceptions, such as the Mamirauá Reserve, in Brazil. However, even in communities where sustainable management is done, there is no consistent, available data concerning the current conservation status of the species, much less concerning the population trends within these communities [10].

Due to the history of disorderly exploitation and the risk of extinction of the Amazonian’s symbol fish *Arapaima gigas*, specialists emphasize the need of using molecular markers to support management strategies and the evaluation of the effectiveness of the *A. gigas*’s conservation programs, highlighting the importance of using these markers for the identification and genetic tracing of marketed specimens [21].

Microsatellite markers, also known as short tandem repeats (STR) or simple sequence repeats (SSR), are DNA sequences consisting of tandemly repeating mononucleotide, dinucleotide, trinucleotide, tetranucleotide and pentanucleotide units, arranged throughout the genomes of most eukaryotic species [22]. They present features such as broad distribution in eukaryotic genomes, which can easily be detected by polymerase chain reaction (PCR), locus-specific nature, co-dominant inheritance, and high mutation rate, being highly polymorphic and hyper-variable [23,24,25]. All these features contribute to the advantages of using microsatellite markers in several research areas – as forensic and population genetics, conservation biology, and for genome mapping in evolutionary and biological scenarios, since these biomarkers allow two or more loci amplifications in a single multiplex PCR reaction.

The multiplex PCR system consists of the simultaneous amplification of various loci, tagged by a distinct fluorescent label, and posterior analysis by capillary electrophoresis in an automated sequencing machine. It is considered a trustworthy technique since the use of capillary electrophoresis with fluorescently labeled primers provides high detection sensibility of the amplified DNA fragments [26]. Several methods are used in microsatellite markers design. However, the majority of these are longstanding and more laborious. With the advances in Next Generation Sequencing (NGS) technologies, increased data became available, allowing a faster, cost-effective, large-scale mining of molecular markers [27,28].

It is urgent to develop new, more accurate and numerous *A. gigas* molecular markers, not only to be applied as a tool for genetic studies, but also to improve the management system, refining the effectiveness of conservation programs, as well as to be used as a tool for the genetic tracing of wild and marketed individuals.

Thus, in order to contribute to the preservation and sustainable management, we developed a broad genomic panel of microsatellite loci potentially amplifiable. Based on this broad panel, we designed an unprecedented multiplex system containing 12 tetranucleotide microsatellite markers, which can be used in populational genetics, conservation biology, and forensic studies of the *A. gigas*.

## 2. Results

### 2.1. Broad panel

A broad amplifiable panel of microsatellites was designed (available at Figshare online data repository – Supplementary Table 1), based on the A. gigas’ genome published by Vialle *et al.* (2018), who obtained a total of 76.91 Gb of high-quality filtered data, such as assembled genome size of 661,278,939 bp, 5,301 scaffolds, scaffold N50 = 668 kb, contig N50 = 51.23 kb, largest scaffold of 5,332,704 (bp), and GC content of 43.18% [29].

We found a set of 95,098 single loci of perfect microsatellites with simple repeat sequences, from the dinucleotide to hexanucleotide classes, in the genome of the *A. gigas* (as observed in the supplementary table 1). The set contained information about the microsatellite motif, the repeat copy number, the initial and final positions, the scaffold and size, the forward and reverse primers, and the expected Polymerase Chain Reaction (PCR) products. The average size of the estimated PCR products was 207 bp. However, we could not design primers for 647 loci, since these sequences were rich in TA repeats, and because the TA motif were at the beginning or at end of the scaffold.

Table 1 presents the abundance of the motifs categorized by class. Here, the dinucleotide class is the most frequent, representing more than 70% of the total microsatellites found in *A. gigas*’s genome. In this class, the TG-repeat motif was the most frequent (15,8). Additionally, we characterized the minisatellites present in 387 loci (Supplementary Table 2).

**Table 1.** Absolute and relative frequency (percentage) of different perfect microsatellite motifs found in the genome of Arapaima gigas.

| Motif class | Count | % |
| --- | --- | --- |
| Dinucleotides | 71,269 | 74.94 |
| Trinucleotides | 13,026 | 13.70 |
| Tetranucleotides | 8,785 | 9.24 |
| Pentanucleotides | 1,107 | 1.16 |
| Hexanucleotides | 911 | 0.96 |
| Total | 95,098 | 100 |

The number of repeats in microsatellites present in the genome of *A. gigas* was quite variable among the 6 classes, presenting an inverse relationship with the size of the motif. The dinucleotide class, for example, presented the largest number of repeats, whereas the class of hexanucleotides presented fewer repeats (Figure 1).

**Fig1.**
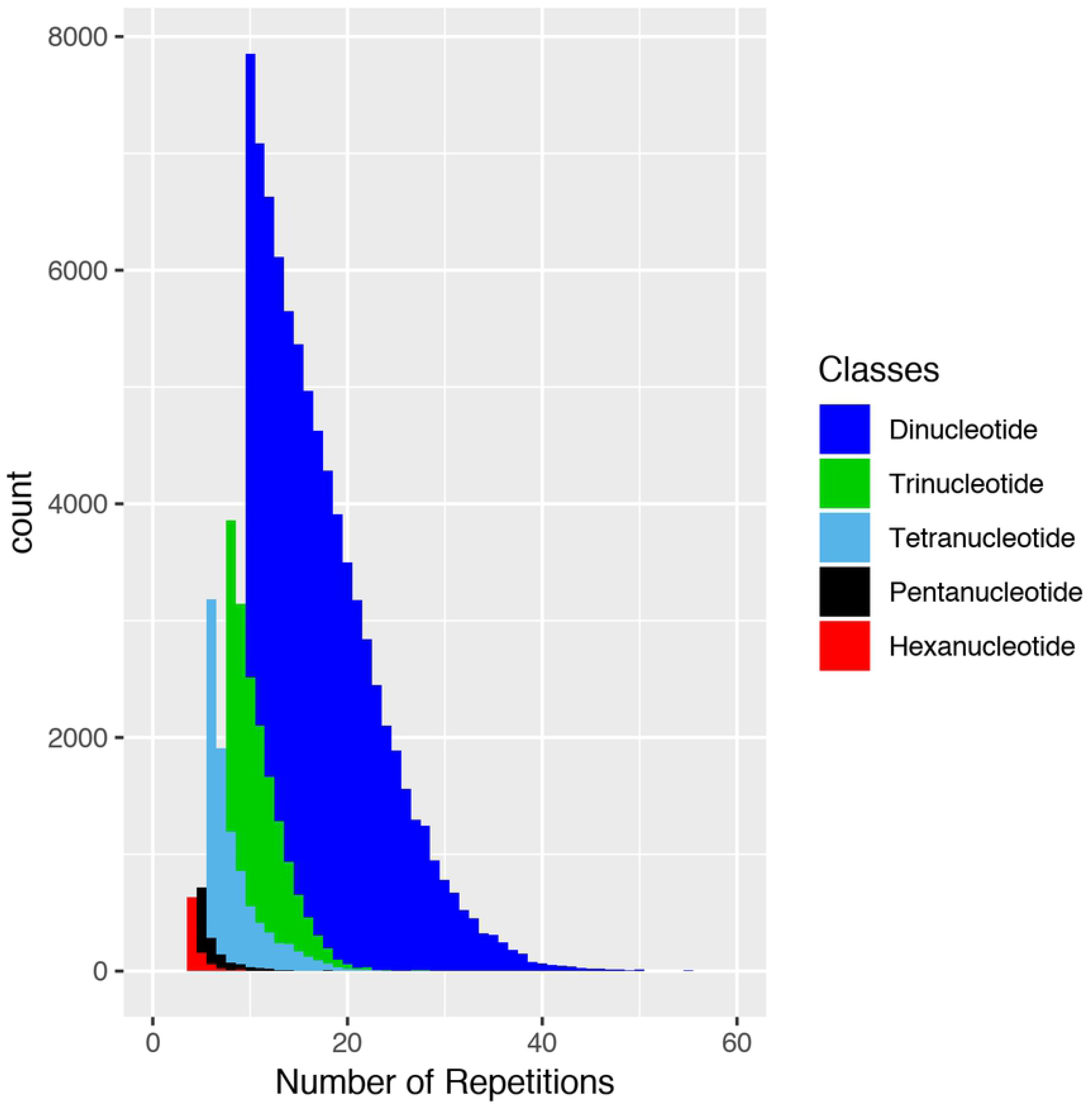
Distribution of the classes of *Arapaima gigas* microsatellites how much to number of repetitions.

The majority of the identified microsatellites were located out of exon and intron sequences, referred here as genomic regions: 68.1% of these were dinucleotides (71,269), 67.8% were trinucleotides (13,026), 67.6% were tetranucleotides (8,785), 67.4 % were pentanucleotides (1,107), and 69.9% were hexanucleotides (911). We observed a few microsatellites located within gene coding regions, 8.3% of which were dinucleotides (5,915), 8.1% were trinucleotides (1,054), 8.3% were tetranucleotides (725), 8.2% were pentanucleotides (91) and 7.8% were hexanucleotides (71) (Figure 2).

**Fig2:**
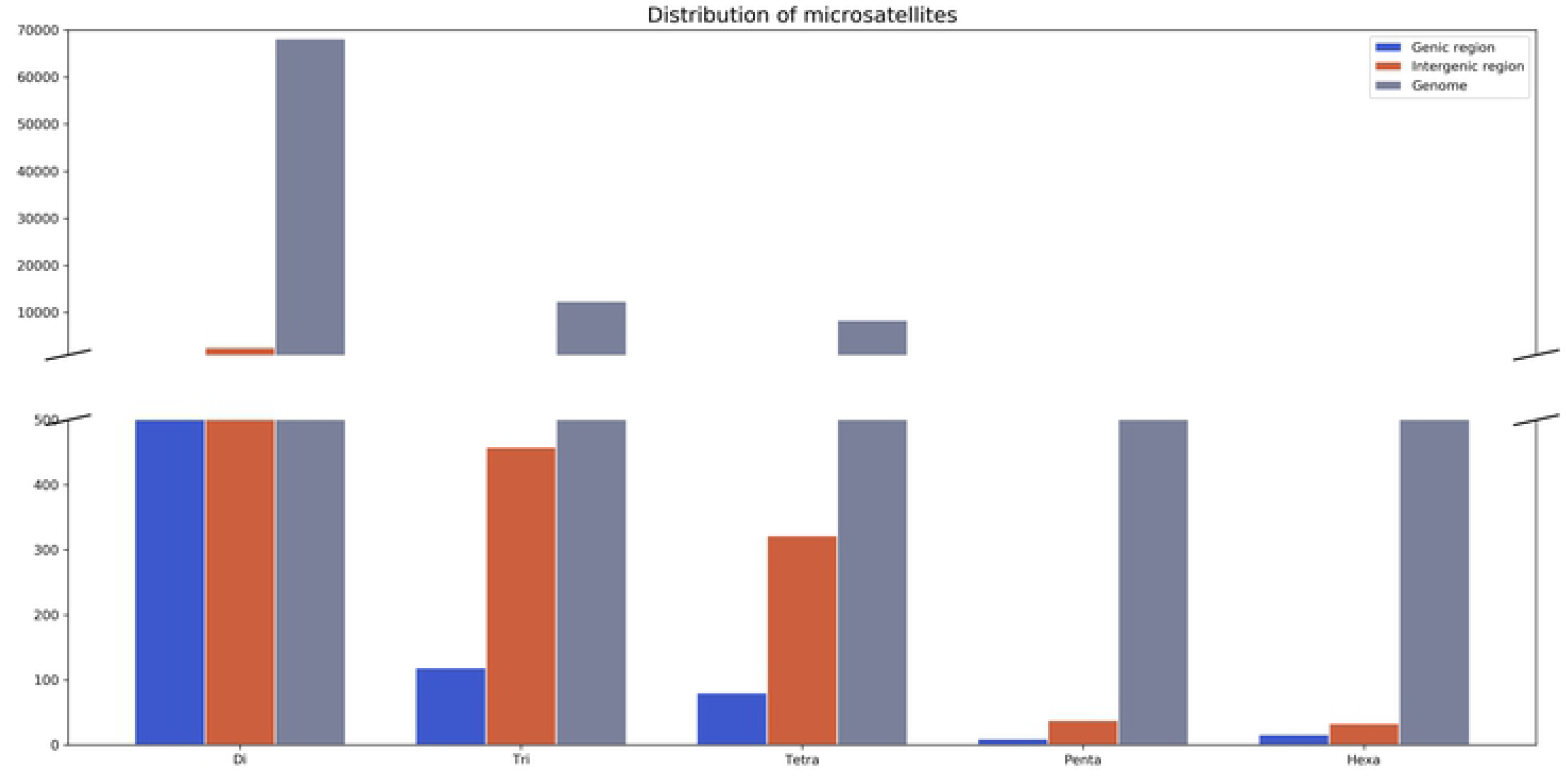
Distribution of microsatellites in genic, intergenic and genome regions in *Arapaima gigas*.

### 2.2. Multiplex system

The broad panel generated in our study allowed the fast selection and validation of 12 tetranucleotide microsatellite loci on a multiplex system (Agig13519, Agig50571, Agig58115, Agig08356, Agig67103, Agig93614, Agig33291, Agig90836, Agig05001, Agig70664, Agig08912 and Agig06409). The multiplex system presented a high resolution, with no overlap between the microsatellite alleles and no artifact peaks (Figure 3). Moreover, we did not detect any genotyping errors attributed to stutter bands, large allele dropouts or null alleles, which are frequent in dinucleotide microsatellites.

**Fig3.**
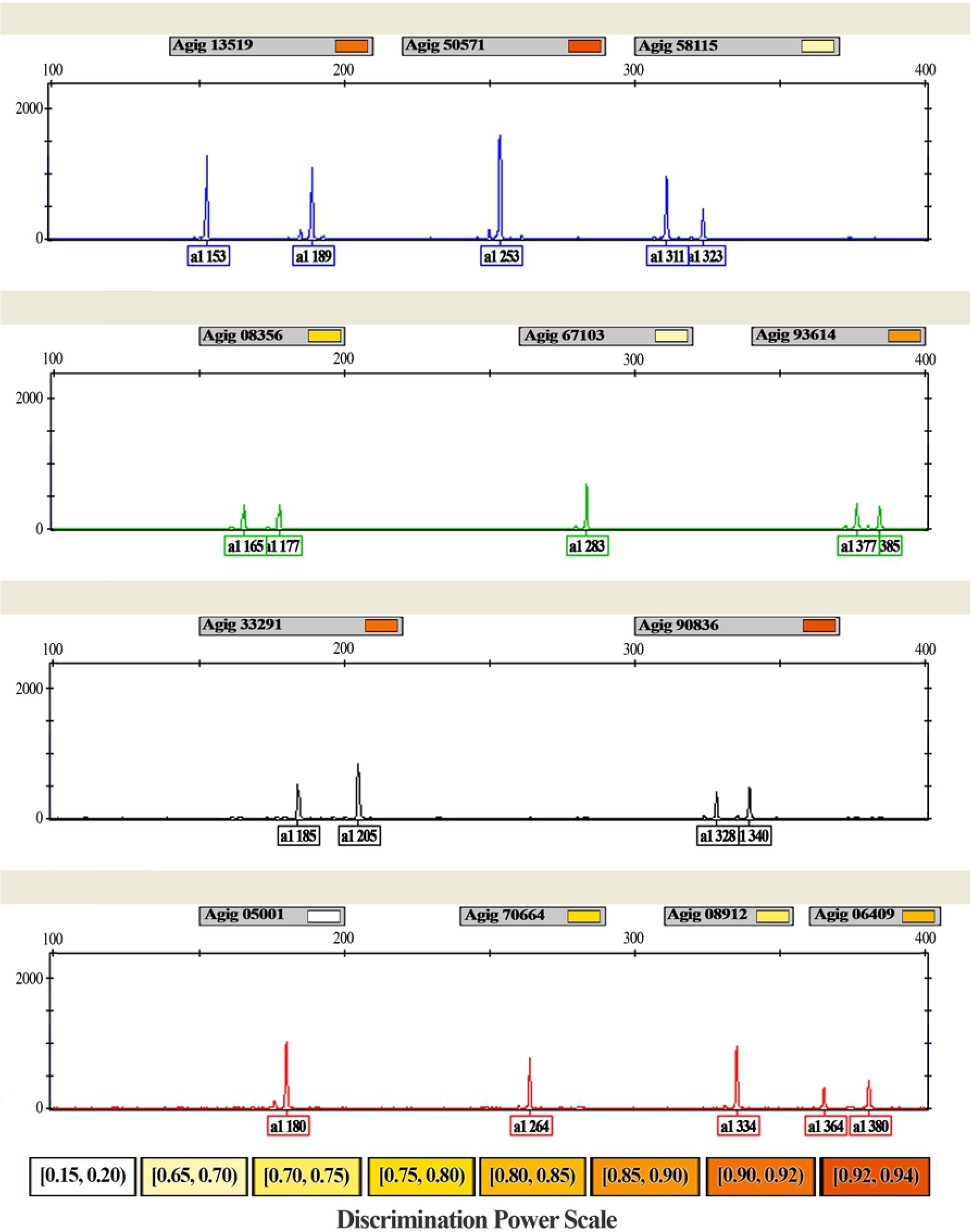
Multiplex PCR panel electrophoretogram for Arapaima gigas showing the allele size range obtained using the ABI 3130 Genetic Analyzer (Applied Biosystems) and the GeneMapper 3.7 (Applied Biosystems) software and discrimination power per locus. Colors were assigned to microsatellite primers labeled with 6-FAM (blue), HEX (green), NED (black), and PET (red) fluorescent dyes.

The panel was used to evaluate a wild population of A. gigas from Santarem (n=30). The obtained data revealed a total of 73 alleles. The average rate of alleles per locus (NA) is 6.08. The observed (HO) and expected (HE) heterozygosity rates range from 0.867 (Agig 33291) to 0.100 (Agig 05001), with an average of 0.59 and 0.64, respectively (Table 2).

**Table 2.**
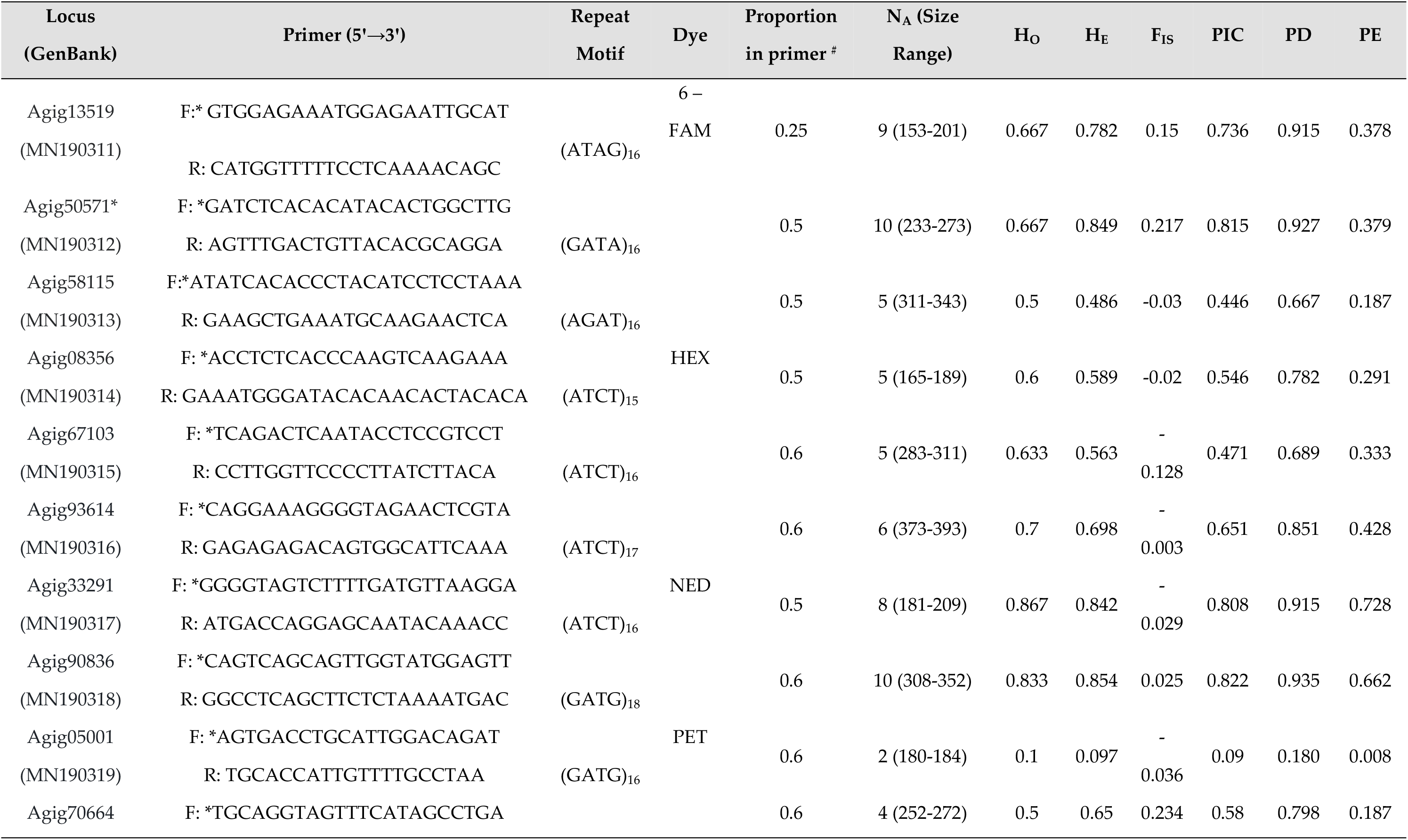
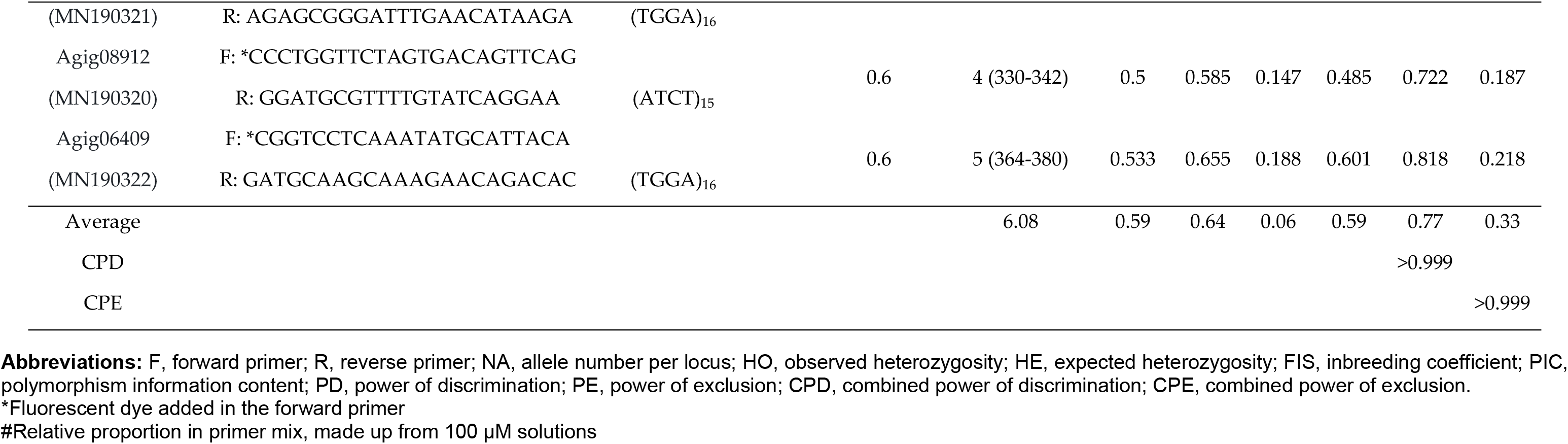
PCR multiplex panel characteristics of 12 microsatellite markers of Arapaima gigas, which was used to genotype 30 individuals from Santarém, a city in the Brazilian Amazon.

There was no significant deviation from Hardy-Weinberg equilibrium (HWE) after Bonferroni correction (p<0.0041). Loci pairs in linkage disequilibrium (LD) were not registered. The average inbreeding coefficient value (FIS) was 0.06.

Forensic parameters investigated for this multiplex system presented average polymorphic informative content (PIC) value of 0.53. The average power of discrimination (PD) was 0.77 and the power of exclusion (PE) was 0.33. The average combined power of discrimination (CPD) and the combined power of exclusion (CPE) for the 12 microsatellite markers investigated was 0.999 and 0.999, respectively (Table 2).

## 3. Discussion

The rapid development of sequencing technologies allowed the obtention of complete genome sequences from an increasing number of species [30], which is an excellent source for identification of microsatellite markers already used in several species [30,31,32,33,34].

In this study, we identified a total of 95,098 microsatellites based on sequences of the complete genome of *A. gigas*, a value close to that found for *Monopterus albus* (99,293) [32] and much larger than what was found for *Brycon orbignyanus* (81,241) [31]. These differences probably are due to the size of the databases, search criteria and bioinformatics tools used to identify the microsatellites [35]. The dinucleotide class was the most abundant, following the same pattern observed for the species mentioned above.

Microsatellite markers with number of repeats among 15-20 tend to be highly polymorphic [36], being the most indicated for population genetic studies. The panel of microsatellite markers developed for *A. gigas* included the dinucleotide, trinucleotide and tetranucleotide classes presenting motifs with repeats within this range (figure 1), however the tetranucleotides markers produced weaker stutter bands and had no artificial multiband patterns [37].

Microsatellite markers are observed in almost all known eukaryotic and prokaryotic genomes, both in coding and noncoding regions [38]. In the *A. gigas* genome, microsatellite markers are more abundant in non-coding sequences, here called genomic regions (figure 2). This corroborates data available in the literature towards the distribution of these markers – which may be explained by the fact that, in promoter regions, the length of microsatellites may influence the transcription activity [39].

It is suggested that microsatellite polymorphisms are associated with the number of loci replications [40]. Microsatellites with higher number of repeats are more prone to mutation/expansion than those with fewer replicates. The correlation between repeat length and microsatellite variability is comprehensible according to replication slippage model, which is a widely accepted mutation mechanism [41]. Another variable could be the microsatellite classification, and their results indicate that tetranucleotide microsatellites have the highest polymorphism rate, while the dinucleotide microsatellites are the least polymorphic [40]. Hence, we emphasize the importance considering the number of repeats and the class of the microsatellite while selecting markers for the accomplishment of scientific studies.

In the last years, several studies aiming to elucidate population genetics and conservation biology of the A. gigas have used dinucleotide microsatellites as molecular markers, as reported by Farias *et al.*, 2003 [42], among other studies [43,44,45,46, 47,48]. However, the differences in quantity and types of microsatellite markers adopted by each study complicate the evaluation of the analyzed indexes among the examined populations, thus the use of a common system for all studies would facilitate the integration of the data for decision making management.

To validate our findings, we developed a multiplex system using the panel of 12 microsatellite markers to evaluate the conservation status of an *A. gigas* population. This tetranucleotide microsatellite markers of the multiplex system have the advantage of being highly polymorphic, more stable and presenting clearer bands than the dinucleotide markers described in literature. Thus, overall, they are becoming the predominant type of microsatellite markers in use [49].

It is important to emphasize that the history of fierce fishing exploitation of the *A. gigas* is a determinant factor for the findings of this study in the population of Santarem. The reduction of natural populations and the decrease in average volume and length of the specimens landed in the Amazon due to overfishing soon lead to the fishing collapse in 1970 [50], occasioning a population bottleneck effect, consequently causing loss of genetic variability, as observed in the values of HE, HO, FIS and NA.

Among the investigated forensic parameters, the values of PIC were considered satisfying accordingly to the scale reported by Botstein *et al.*, 1980 [51]. The CPD and CPE values of this panel allow the distinction of one individual in one billion, which is similar to other multiplex genotyping systems. For instance, the system reported by Hamoy *et al.*, 2012 [52] was used in conservation studies and in the aquaculture support.

Population genetics data are undoubtedly the most important component of the baseline of any conservation and management plan [53]. The broad microsatellite panel developed for *A. gigas* opens perspectives for studies in as many fields as bioinformatics, ecology, genetics, evolution, and comparative studies among species [31]. And the multiplex system designed for *A. gigas* allows an accurate, faster, cost-effective, and affordable genotyping for low-income laboratories and conservation studies focusing on this species.

The set of 12 loci was able to measure the genetic variability of the investigated population, providing high statistic power data, sufficient for determining kinship patterns for population attribution tests. Therefore, this multiplex system can represent a valuable and powerful tool for small and large-scale studies in the areas of forensic/conservation biology and population genetics, furthermore, it can also be used as a tool for the management of wild and cultivated populations of *A. gigas*.

## 4. Materials and Methods

### 4.1. Data mining

The development of this study was based on the complete genome of *Arapaima gigas*, published by Vialle *et al.*, 2018 [29], available from the National Center for Biotechnology Information (NCBI) database (https://www.ncbi.nlm.nih.gov/assembly), ID: 12404.

### 4.2. Bioinformatics analysis

Simple Sequence Repeat Identification Tool (SSRIT) [36] was used to find all simple sequence repeats (SSRs). We changed the default parameters related to motif-repeat in the source code of SSRIT, which was made in Perl, to specify motif length and the minimum number of repeats (Table 3). The other attributes remained with their default values.

**Table 3.**
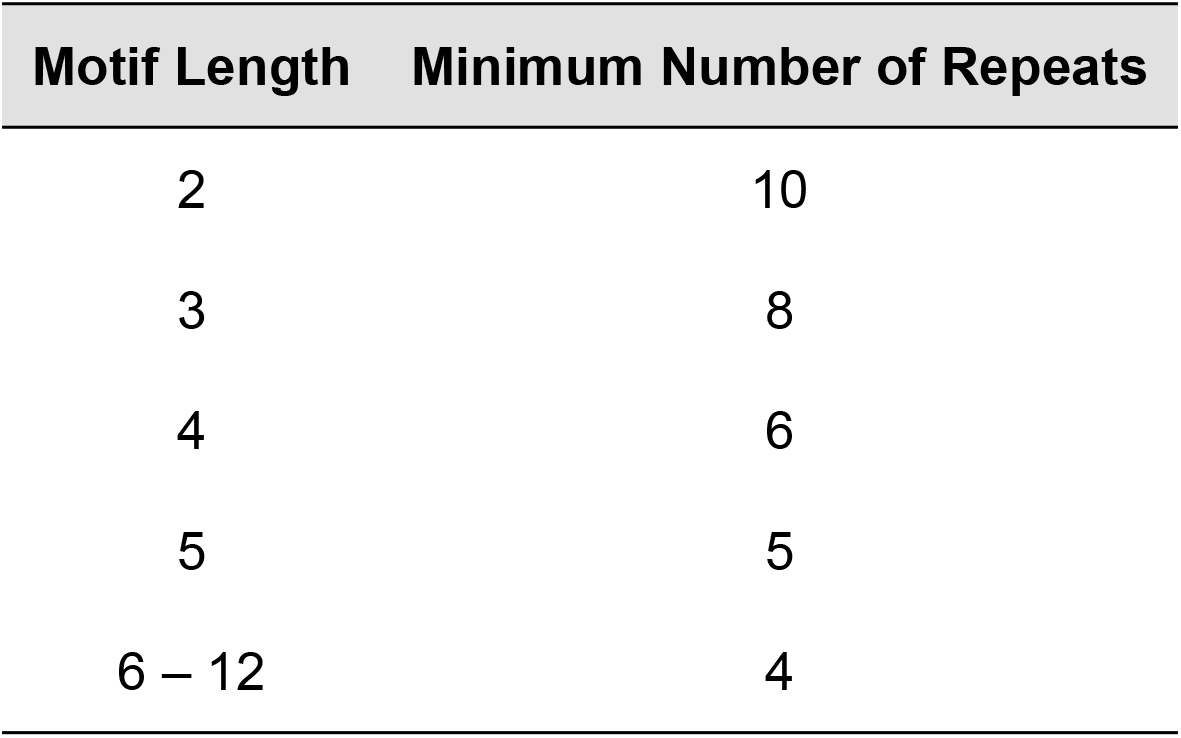
Parameters Modified in SSRIT Analysis.

| Motif Length | Minimum Number of Repeats |
| --- | --- |
| 2 | 10 |
| 3 | 8 |
| 4 | 6 |
| 5 | 5 |
| 6 – 12 | 4 |

Microsatellites sequences counted from all samples generated by the SSRIT were processed by in-house scripts in Python (v. 3.7.2) to insert the flanking regions in the 5’ and 3’ portions of DNA sequence, and transform them in a fasta file. After, the sequences which exhibited microsatellites were submitted to Primer 3 (v. 4.1.0), whose premise is design PCR primers from DNA sequences. In Primer 3, the default parameters of primer length, (|Primer Opt Size = 20|, |Primer Min Size = 18|, and |Primer Max Size = 23|) were modified to 24, 20, and 28, respectively. The parameters of primer product size range was modified for 50-1200 nucleotides.

The microsatellites were individually filtered by abundance and number of repetitions per class, and subsequently mapped in regions of intron (intergenic region) and regions of exon (exonic regions). The ones which are not present in any of the previous regions were classified as belonging to the genome. The development of all figures was made using Matplotlib (v. 3.0.1), which is a Python 2D plotting library.

### 4.3. Validation of the multiplex microsatellite system

- Sample Collection and DNA Extraction

This research was approved by the Ethics Committee on Animal Use (CEUA) of the Federal Rural University of Amazon - UFRA, protocol number 055/2017 (CEUA) – 23084.017501/2017-02 (UFRA).

We collected 2 g of muscle tissue of 30 specimens of *A. gigas* from a fishing vessel landed at the city of Santarem, localized in the Low Amazon mesoregion, Brazil. The samples were preserved in 70% ethanol and posteriorly stored in −20 °C.

Total genomic DNA was extracted from the digested tissue in proteinase K solution/ Sodium Dodecyl Sulfate (SDS) and purified in Phenol/Chloroform, followed by precipitation in Isopropanol [54]. The DNA concentration was measured in the NanoDrop ND1000 spectrophotometer (Thermo Scientific).

- Selection of Microsatellites and Polymorphism Testing

After assembling the broad microsatellite panel for *A. gigas*, a total of 12 microsatellite markers were selected to compose the multiplex system, according to the following criteria: tetranucleotide microsatellites, with at least 15 repeats and at maximum 20 repeats, outside of coding regions. The polymorphisms of the 12 selected loci were tested in 7% polyacrylamide gel electrophoresis, stained with silver nitrate.

- Primer Testing and Genotyping

The possibility of formation of secondary structures among the primers was tested in AutoDimer Sofware [55]. A PCR reaction consisting of the simultaneous amplification of 12 markers was standardized to a final volume of 9.5 μL, using 5.0 μL 2X Qiagen Multiplex PCR Master Mix (Qiagen),S 1.0 μL of Q-Solution (Qiagen), 2.0 μL of H2O, 0.5 μL of primer mix, and 1.0 μL of genomic DNA. The relative proportion of each primer in the primer mix (made up of 100 μM solutions) is listed in Table 1. The reactions were optimized to amplify 5 ng of genomic DNA.

Amplification reactions were performed in a Veriti thermocycler (Applied Biosystems). The thermocycling conditions were: initial denaturation at 95°C for 15 min, followed by 10 cycles at 94°C for 30 s, 60°C for 90 s, and 72°C for 60 s; 20 cycles at 94°C for 30 s, 58°C for 90 s, and 72°C for 60 s, and a final extension at 72°C for 60 min, 10° for min. One microliter of amplification was mixed with 8.5 μL Hi-Di deionized formamide (Applied Biosystems), and 0.5 μL GeneScan 500 LIZ (Applied Biosystems) as a molecular weight standard. The final product was analyzed using an ABI 3130 Genetic Analyzer (Applied Biosystems). The determination of fragment size and allele designation was done with the GeneMapper 3.7 software (Applied Biosystems).

### 4.4. Statistical Analysis

The dataset was checked for genotyping errors and null alleles using Micro-Checker [56]. We analyzed the genetic variability using the allele number per locus (N_A_), the observed (H_O_) and expected (H_E_) heterozygosity indexes, and the deviation from Hardy-Weinberg equilibrium (HWE), using Arlequin 3.5.1.257, followed by Bonferroni’s correction [58]. The same program was used to determine the proportion of locus pairs in linkage disequilibrium (LD).

The inbreeding coefficient (F_IS_) was calculated in GENEPOP [59]. The polymorphism information content (PIC), the power of discrimination (PD), and the power of exclusion (PE) for all markers using the forensic statistic tool FORSTAT [60].

## Supplementary Materials

Supplementary materials can be found Supplementary Table 1 - https://doi.org/10.6084/m9.figshare.8088533) and Supplementary Table 02 - https://doi.org/10.6084/m9.figshare.8088629).

## Author Contributions

Conceptualization, P.F.-G.and J.A.; methodology, P.F.-G, D.M., H.P., F.M., R.C., J.S. and C.S; software, H.P., F.M., R.C., J.S.; validation, P.F.-G. and C.S; formal analysis, P.F.-G., R.C. and J.S.;investigation, P.F.-G.and J.A.; resources, M.R.; I.H. and S.S.; data curation, P.F.-G., J.A, D.M. and G.C.; writing—original draft preparation, P.F.-G; writing—review and editing, P.F.-G., J.A. and G.C.; visualization, S.S., M.R. and I.H.; supervision, S.S.; project administration, S.S. and I.H.; funding acquisition, M.R.; I.H. and S.S.; All authors have read and agreed to the published version of the manuscript.

## Acknowledgments

We thank the Federal University of Pará and Rural Federal University of the Amazon for all the support.

## Abbreviations

CITES: International Convention on the Trade in Endangered Species
IUCN: International Union for Conservation of Nature’s
STR: Short Tandem Repeats
SSR: Simple Sequence Repeats
PCR: Polymerase Chain Reaction
NGS: Next Generation Sequencing

